# GluN2A-mediated currents and calcium signal in human iPSC-derived neurons

**DOI:** 10.1101/2025.09.26.678211

**Authors:** Sergio Escamilla, Carlos Avilés-Granados, Francisco Andrés Peralta, Ana V Paternain, María-Ángeles Cortés-Gómez, Henrik Zetterberg, Elvira de la Peña, Federico Salas-Lucia, Javier Sáez-Valero, Inmaculada Cuchillo-Ibáñez

## Abstract

Gene expression data indicate that during human brain development, neurons change the NMDA receptor (NMDAR) subunit composition to modulate their function, favouring the GluN2A subunit over GluN2B - a hallmark of neuronal maturation. However, evidence supporting this phenomenon in human iPSC-derived neurons remains elusive. Here, using two differentiation methods in parallel (BrainPhys Neuronal Medium, BPM, and Neural Maintenance Medium, NMM), we provide evidence of increased synaptic localization of NMDARs during neuronal maturation and that GluN2A subunit is crucial for the NMDA physiological function-inducing inward currents and calcium entrance at 60 days of differentiation. Calcium responses to specific agonists, particularly NMDA, were elevated in cells cultured under BPM conditions. This is likely attributable to their more mature neuronal phenotype and the RNA-seq-identified upregulation of genes involved in intracellular calcium signaling proteins. Our results offer insight into how glutamate receptor subunits mature during brain development, delineating approaches to study NMDAR activity in health and disease.

**SUMMARY:** This study shows that GluN2A subunit is essential for proper NMDAR function in cultured human neurons, evidenced by changes in intracellular calcium and ionic currents after specific agonist exposure. This places GluN2A at the crossroads of developmental and degenerative disease

## INTRODUCTION

During brain development, the correct maturation of excitatory glutamatergic neurons is essential for the function of this organ later in life (Greig et al., 2013). As glutamatergic neurons mature, they progressively establish synaptic connections with other neurons (Prince et al., 2024; Wallace & Pollen, 2024) using different glutamate receptors. Among these, N-methyl-D-aspartate receptors (NMDARs) play a key role in their maturation (Hunt & Castillo, 2012; Ikonomidou et al., 1999; Lee et al., 2005). NMDARs are constituted by four subunits; two obligatory GluN1, and the composition of the other two depends on the different brain areas and developmental stages (Sanz-Clemente et al., 2013). In regions of the human brain such as the cerebral cortex, hippocampus and cerebellum, the alternative subunits GluN2B and GluN2A are the most abundant (Kornhuber et al., 1989; Paoletti et al., 2013; Sanz-Clemente et al., 2013; Scherzer et al., 1998; van der Aart et al., 2019).

However, the abundance of GluN2B and GluN2A subunits changes across brain development. Seminal studies (Bagasrawala et al., 2017; Bar-Shira et al., 2015; Jantzie et al., 2015; Pegasiou et al., 2020) looking at the GluN2B and GluN2A mRNA expression in the cerebral cortex, hippocampus and cerebellum of human fetuses describe a switch in the abundance of these subunits at different gestational weeks. At early gestational weeks, GluN2B subunit expression is high, whilst GluN2A expression is low. This changes during the final stages of prenatal development, when the GluN2A subunit becomes the most abundant. Notably, the GluN2B-GluN2A switch during brain development is conserved across vertebrates, including amphibians, birds, and mammals (Dumas, 2005), indicating a relevant role in the development of this organ.

Expression of both, GluN2B and GluN2A subunits have a high relevance for proper and healthy brain function. Several population studies have shown a link between genetic variants in the genes encoding these subunits (*GRIN2B* and *GRIN2A*) and a higher prevalence of several neurodevelopmental disorders, including autism spectrum disorders, schizophrenia, epilepsy and intellectual disability (Harrison & Bannerman, 2023; Mangano et al., 2022; Reutlinger et al., 2010). Further evidence comes from studies in chimeric mouse models that express human proteins associated with neurological and motor disorders. These studies describe changes in the GluN2B-GluN2A switch, suggesting that alterations in this process may contribute to the pathophysiology of neurological conditions (Maltese et al., 2018; Sinclair et al., 2016).

Studies in rodents have shown that the GluN2B-GluN2A switch represents a key process in neuronal maturation and the refinement of neuronal circuits (Barria & Malinow, 2002; Flint et al., 1997; Sanz-Clemente et al., 2013; Sheng et al., 1994). This becomes clear in mice deprived of visual input during development, that leads to the absence of the GluN2B-GluN2A switch in the visual cortex (Philpot et al., 2001), and strongly suggests that the GluN2B-GluN2A switch requires synaptic input to take place (Yashiro & Philpot, 2008). In addition, studies in mice show that the GluN2B-GluN2A switch is implicated in synaptic plasticity (Baez et al., 2018; Bellone & Nicoll, 2007; Franchini et al., 2019), as illustrated by changes in electrophysiological properties (e.g., increased excitatory postsynaptic current amplitude and reduced decay time) that indicate this switch enhances synaptic efficiency and maturation (Kirson & Yaari, 1996; Thomas et al., 2006). Collectively, these findings place the GluN2B-GluN2A switch central to the mechanisms underlying neuronal maturation (Bar-Shira et al., 2015; Law et al., 2003; Stocca & Vicini, 1998; Tovar & Westbrook, 1999). However, since access to human fetal brain tissue is limited, the contribution of the GluN2B-GluN2A switch to the electrophysiological properties and integrated neural response of human neurons remains largely unexplored.

Here, we leverage two different protocols for generating neurons from stem cells to study the appearance of the GluN2B-GluN2A switch. Together, our findings demonstrate that human iPSC-derived neurons exhibit significant GluN2A-dependent ionic currents and calcium signaling, supporting that the GluN2B-GluN2A switch not only takes place but also influences the functional properties of human iPSC-derived glutamatergic neurons. Our results provide insight into how glutamate receptor subunits act during brain development, laying the groundwork for delineating the mechanisms of action of these components in developmental, behavioral and degenerative diseases.

## RESULTS

### Characterization of iPSC-derived neural cultures

We used one iPSC line that formed compact colonies with smooth edges and expressed pluripotency markers, with >80% of iPSCs expressing both OCT4 and Nanog (**Supplemental Figure 1A, B, C**). iPSCs were treated for 18-21 days with inhibitors of the SMAD signaling (GSK3, SMAD and NOTCH) (Chambers et al., 2009) to obtain a 2D culture of neural precursor cells (NPCs) expressing PAX6 and Nestin (**Supplemental Figure 1D).** NPCs were further differentiated into neural cultures. We employed two different culture protocols to induce neuronal maturation (details in the methods section). In one protocol, we used BrainPhys Neuronal Medium (BPM conditions), that favors neural maturation and results in cultures with many astrocytes (von Bartheld et al., 2016); in the second, we used Neural Maintenance Medium (NMM conditions), that promotes less neuronal maturation and less astrocyte differentiation (Shi et al., 2012) (**Figure 1A**) when compared to BPM.

**Figure 1.**
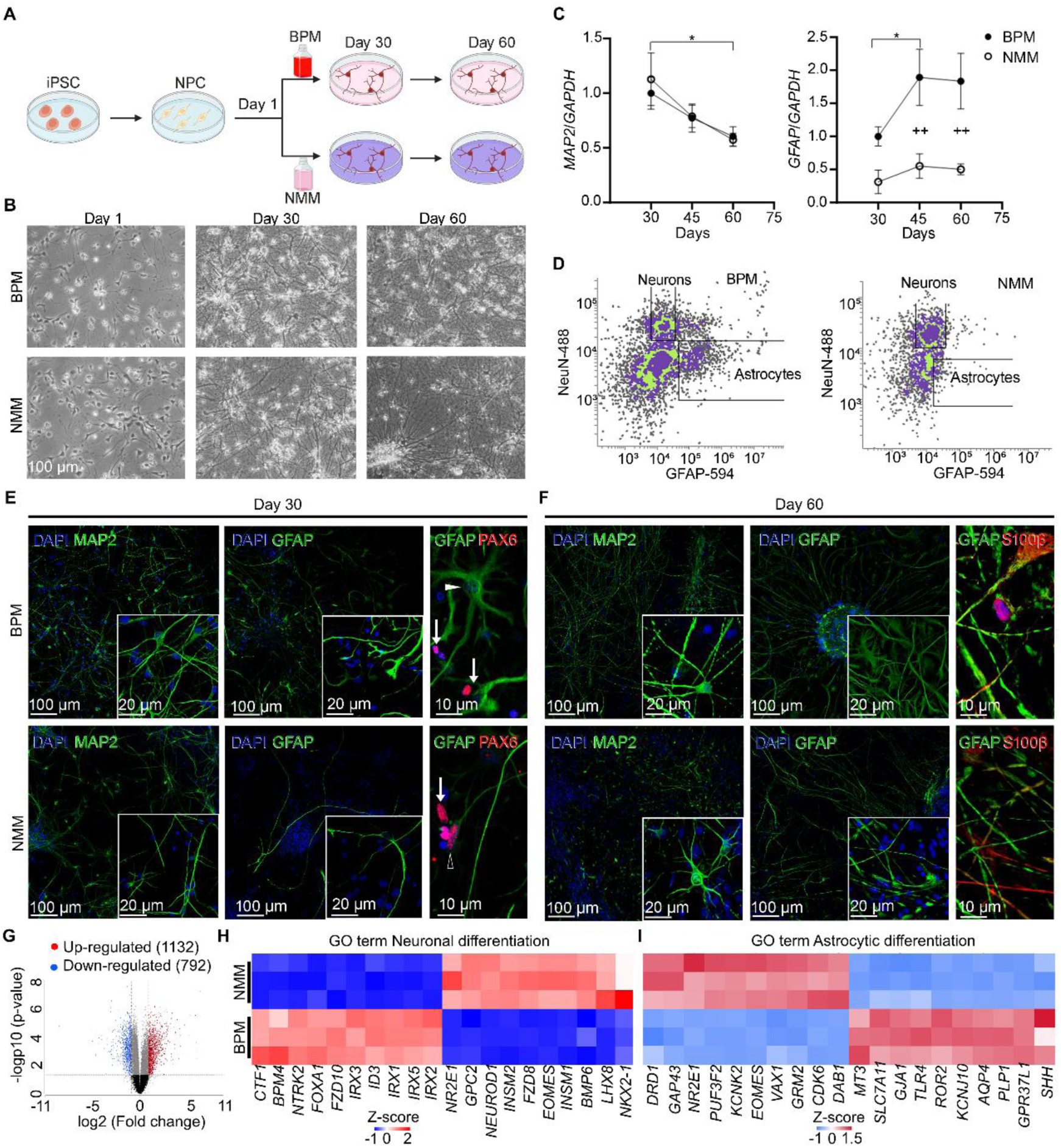
Characterization of iPSC-derived neural cultures. **A)** Schema of the iPSCs neurodifferentiation using either BPM or NMM. **B)** Brightfield microscope images showing iPSC-derived neural cultures using either BPM or NMM at different time points of differentiation after NPCs platting. **C)** Relative mRNA levels of *MAP2* and *GFAP* respect to *GAPDH* at different time points during the process of neurodifferentiation. For *MAP2*, “*” indicates statistical differences between time points for BPM and NMM conditions; for *GFAP*, “*” indicates statistical differences between time points for BPM only, and “+” indicates statistical differences between BPM and NMM. Two-tailed Student’s *t* test \**p* < 0.05, ++*p* < 0.01. **D)** Dispersion plots from FACS experiment of cultures in either BPM or NMM conditions showing cell singlets; neurons are immunolabeled with NeuN-488 and astrocytes with GFAP-594. To ensure high-confidence data, only singlets with strong signals from both fluorophores were included in the statistical analysis (indicated within the black boxes). **E, F)** Immunofluorescence images of neural cultures in either BPM or NMM conditions at Day 30 (E) and Day 60 (F), stained for endogenous MAP2 (neuronal marker) and GFAP (astrocyte marker). The white arrowhead shows a canonical star-shaped GFAP+ astrocyte (immunopositive for GFAP); white arrows show PAX6+/GFAP-cells; empty arrowhead indicates PAX6+/GFAP+ cells (only present in cells cultured in NMM). At Day 60 many astrocytes were GFAP+/S100β+ in both conditions. **G)** Volcano plot showing the distribution of differentially expressed genes between neural cultures (BPM *vs* NMM) at Day 60 of neurodifferentiation. **H, I)** Heatmap extracted from the RNA-seq data showing genes associated with the GO cluster “Neuronal differentiation” (H) and “Astrocytic differentiation” (I). Thresholds of differentially expressed genes: *p*-value < 0.05, Average Log2 Fold-Change of 1.5 in the Partek Flow platform. Expression values are mean ± SD of n= 3-6 RNA samples, each of which consists of 2 pooled 12-well plates of iPSC-derived neurons.

Once NPCs were plated (day 1; D1) in the two culture conditions, we monitored them throughout their neural differentiation until D60 (**Figure 1B**). During this period, we evaluated the presence of neurons and astrocytes. We observed a decrease in the mRNA levels of the neuronal gene *MAP2* in both conditions, but an increase in the astrocytic gene *GFAP* only in cells cultured with BPM conditions (**Figure 1C**). Next, we collected about 5x10^6^ neural cells at D60 and using fluorescence-activated cell sorting (FACS), we found that neural cells cultured with BPM exhibited a 1:1 ratio of NeuN-positive neurons to GFAP-positive astrocytes, while this ratio was 8:1 in neural cells cultured with NMM (**Figure 1D, Supplemental Figure 1E**). We further evaluated the presence of neurons and astrocytes at D30 and D60 using immunofluorescence. During this period, we observed an increase in MAP2-positive neurons and GFAP-positive astrocytes (**Figure 1E, F**) in both conditions but, already from D30, the cells exhibited a more complex morphology when cultured with BPM. In agreement with the FACS data, the number of neurons was similar between conditions, but the number of GFAP-positive astrocytes was higher in the neural cultures in BPM conditions than in NMM. To confirm that these cells were indeed astrocytes, we used S100β as a marker of mature astrocytes. As expected, at D60, most of GFAP-positive astrocytes were also positive for S100β.

### Transcriptome analysis shows that iPSC-derived neurons cultured in BPM are more mature than those cultured in NMM

To assess the degree of maturation of iPSC-derived neurons between conditions, we performed bulk RNA-seq on neural cells at D60; the datasets clustered separately in a principal component plot (**Supplementary Figure 1F**), indicating significant differences in their transcriptomes.

In fact, neural cells cultured in BPM resulted in 1924 differentially expressed genes (1132 upregulated; fold-change ≤2 to ≥2) compared to the neural cells cultured in NMM (**Figure 1G; Supplementary Table 1**). Remarkably, 3 out of the top 6 up-regulated genes were within the *DLK1*-*DIO3* locus (*DLK1*, *RLT1* and *DIO3* genes), which is involved in neurogenesis and neuronal maturation (Surmacz et al., 2012; Yen et al., 2018). Other top-up-regulated genes included *BMP4*, which encodes a secretory protein that stops proliferation of NPCs and promotes neurodifferentiation (Jensen et al., 2021). Mutations in *DLK1* and *BMP4* are associated with neurodevelopmental delay and brain malformations (Bakrania et al., 2008; Ioannides et al., 2014).

To study the biological interpretation of these transcriptomic differences, we identified 12 gene sets with enrichment scores higher than 4, all related to neuron differentiation/maturation (**Supplementary Figure 1G; Supplementary Table 2**). Examples of these gene sets are: “neuron differentiation”, “neuron spine”, “neuron fate commitment”, “neuron action potential” and “forebrain neuron fate commitment”. Within the gene set “neuron differentiation”, the top 10 up-regulated genes (**Figure 1H; Supplementary Table 3**) include *NTRK2*, a transmembrane tyrosine kinase receptor that promotes the differentiation of NPCs into neurons (Roussel-Gervais et al., 2023). Other upregulated genes included the *IRX* family, which have been involved in several phases of brain development in Xenopus (de la Calle-Mustienes et al., 2002; Gómez-Skarmeta et al., 1998), zebrafish (Cheng et al., 2007) and mice (Dou et al., 2021); *FOXA1*, a transcription factor involved in neuronal differentiation in mice (Dong et al., 2012), and *PROX1*, a member of the Wnt signaling cascade involved in neurogenesis in mice (Karalay et al., 2011; Lavado et al., 2010). As expected, the “pathway enrichment analysis” found an enrichment in “Glutamatergic synapse” with a score higher than 100 (**Supplementary Table 4**).

We also identified 3 gene sets with enrichment scores higher than 4 related to “astrocyte development”, “astrocyte differentiation” and “astrocyte projection” (**Supplementary Figure 1H**). The upregulated genes in neural cells cultured in BPM include *AQP4,* which encodes a water channel critical for astrocyte function (Salman et al., 2022), and *ROR, GAP43* and *TLR4,* all involved in astrogliosis and inflammation (Hung et al., 2016; Liu et al., 2022) (**Figure 1I; Supplementary Table 1**).

Altogether, these findings suggest that the iPSC-derived neural cells cultured in BPM conditions may contain more mature glutamatergic synapses and a more physiological neuron-to-astrocyte ratio than those cultured in NMM.

### The maturation of glutamatergic neurons leads to an increase in synaptic NMDARs

A key feature of neuronal maturation is the insertion of NMDARs at synaptic membranes (Jiang et al., 2011). However, it is unclear when the NMDARs begin to be present in human neurons along development (Bagasrawala et al., 2017), and this process has not been addressed in iPSC-derived neurons either (Petralia et al., 2010; Zhang et al., 2022). Studies using mouse primary neurons described that after 7 days *in vitro*, only ∼10% of NMDARs localize at the synapse, and this increased to ∼50% after two weeks (Gladding & Raymond, 2011; Groc et al., 2006; Ivanov et al., 2006; Rosenmund et al., 1995; Tovar & Westbrook, 1999, 2002).

We evaluated whether the presence of NMDARs at synapses increases over time in the iPSC-derived neurons. To do so, we measured at D30 and D60 the colocalization of the common subunit of all NMDARs, GluN1, with the presynaptic protein Synaptophysin (**Figure 2A, B, C, D**). We found that the colocalization significantly increased from D30 to D60, indicating an increment of synaptic NMDARs in both conditions, which was significantly higher in neurons cultured in BPM than in NMM at D60 (**Figure 2E**).

**Figure 2.**
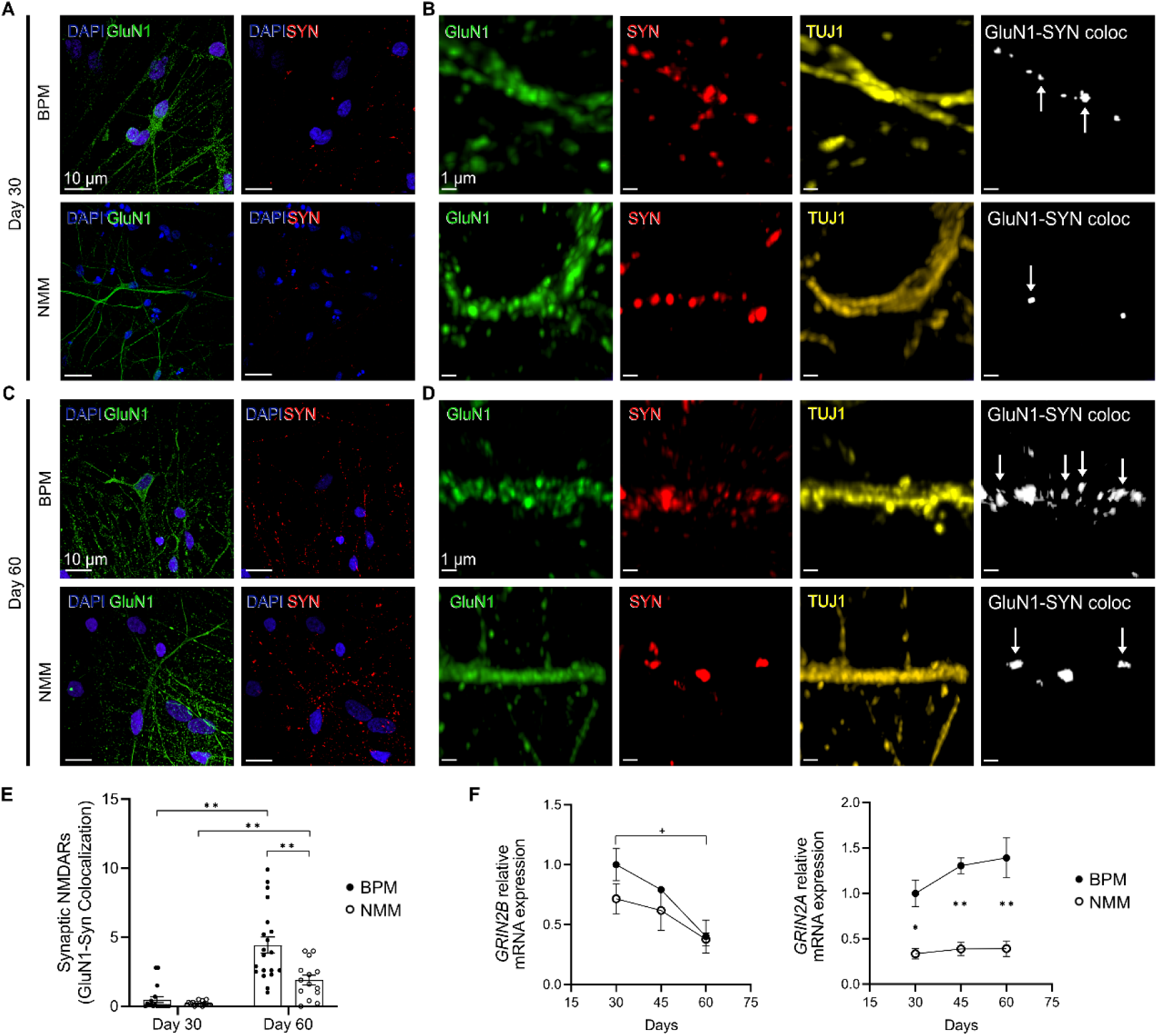
The maturation of the iPSC-derived neurons is accompanied by an increase in the synaptic location of NMDARs and expression of *GRIN2A*. **(A,B)** Immunofluorescence images showing immunopositive cells for GluN1, Synaptophysin and DAPI in neural cultures at Day 30 of neurodifferentiation using either BPM or NMM conditions (A) at low magnification (40x) and (B) at high magnification (40x plus digital zoom). In B, immunopositive cells for TUJ1 and GluN1-Synaptophysin colocalization channel (white arrows). **C, D)** Same as A, B) at Day 60 of neurodifferentiation. **E)** Quantification of the GluN1-Synaptophysin colocalization through Mander’s coefficient at Day 30 and Day 60. **F)** Relative mRNA *GRIN2A and GRIN2B expression*; values were normalized respect to *GAPDH* mRNA levels. Expression values are mean ± SEM of n= 3-6 RNA samples, each sample consists of 2 pooled 12-well plates of neural cultures; “*” indicates statistical differences between BPM and NMM conditions; “+” indicates statistical differences between time points, only for BPM; Student’s *t* test, \**p* < 0.01, \*\**p* < 0.001, +*p* < 0.01.

Neuronal maturation is also characterized by a progressive increase in the expression of the NMDAR subunit GluN2A, which is encoded by the gene *GRIN2A*, and a decrease in the expression of GluN2B, encoded by the gene *GRIN2B*. Although bulk RNA-seq did not show differences at D60 in *GRIN2A* or *GRIN2B*, we tested whether this switch could occur along the maturation by RT-qPCR at different time points. We observed a significant decrease in *GRIN2B* expression from D30 to D60, in both conditions, and a tendency in *GRIN2A* expression to increase along maturation particularly in BPM conditions, that was significantly higher at all-times respect to NMM (**Figure 2F**). This suggests a transcriptomic switch in NMDAR subunits detectable when using targeted RT-qPCR in BPM conditions, which is accompanied by a stronger neuronal differentiation in terms of synaptic NMDAR location.

### GluN2A drives ionic currents in human glutamatergic neurons

The presence of NMDARs at the synapses of iPSC-derived neurons suggests that glutamate can elicit NMDAR-dependent ionic currents in these cells. To test this, we performed whole-cell patch-clamp recordings on iPSC-derived neurons at D60. Prior, we examined whether the iPSC-derived neurons exhibit spontaneous synaptic activity. Under conditions of absence of external input, the cells cultured in BPM showed a higher spontaneous synaptic activity than the ones cultured in NMM (**Figure 3A**).

**Figure 3.**
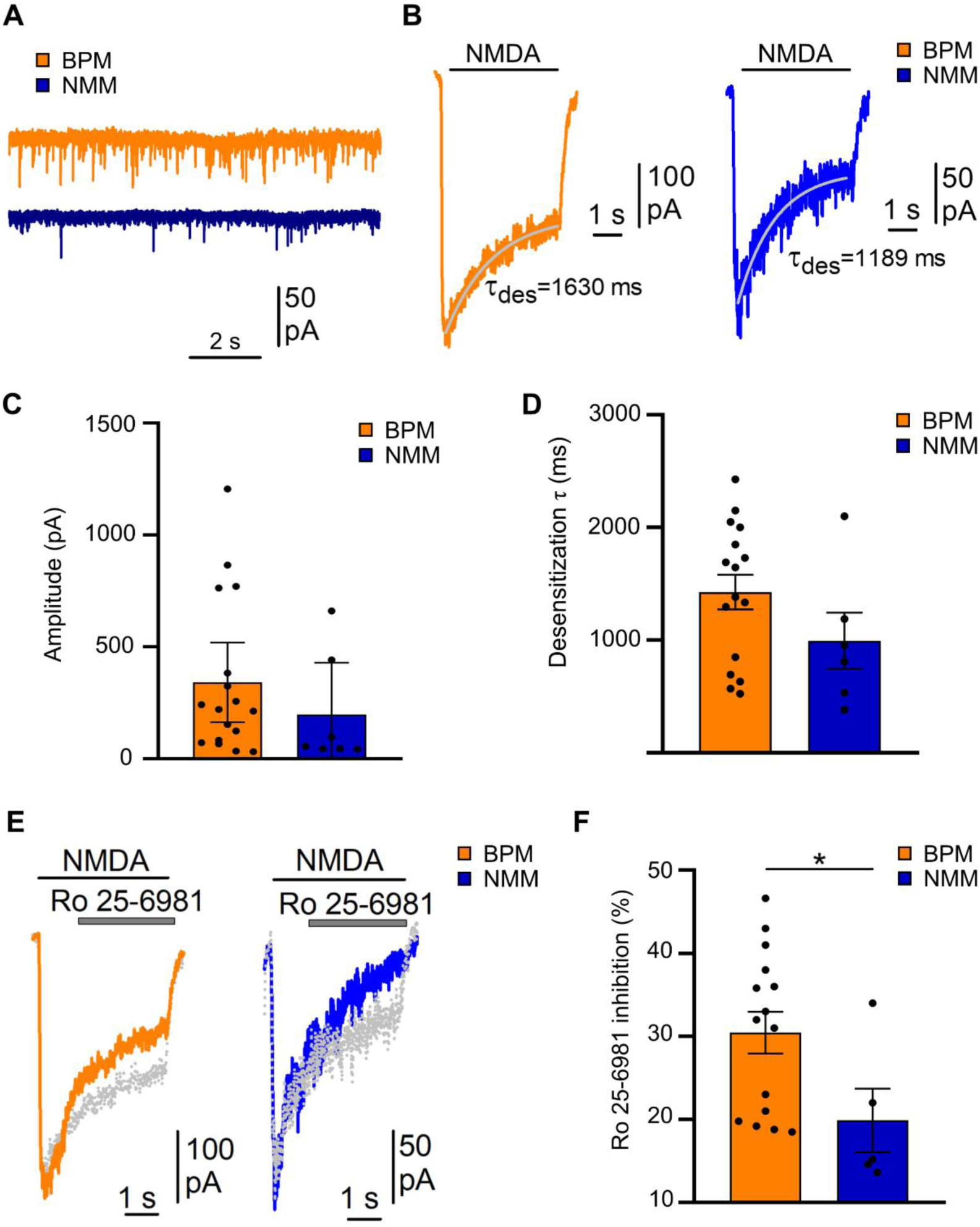
Electrophysiological measurements of NMDA-dependent currents in iPSC-derived neurons in BPM and NMM conditions after 60 days in culture. **A)** Spontaneous activity recorded as excitatory postsynaptic currents in neurons cultured in BPM and NMM conditions. **B)** Representative membrane current upon addition of 100 μM NMDA with 10 μM Glycine in neurons cultured in BPM and NMM. The superimposed grey line illustrates a single exponential fitting the desensitization process with the indicated τ. **C, D)** Quantification of the NMDA-induced current amplitudes (C, n= 17 for BPM, n= 7 for NMM; *p*= 0.2094, Mann-Whitney test) and desensitization τ (D, n= 16 for BPM, n= 6 for NMM, *p*= 0.1775, Mann-Whitney test). **E)** Same as B), upon addition of 1 μM Ro 25-6981 for inhibiting GluN2B containing receptors. Superimposed grey traces are the responses without blocker added. **F)** Percentage of Ro 25-6981 inhibition respect to NMDA-induced current (n= 15 for BPM, n= 5 for NMM, \**p* < 0.05, Two-tailed Student’s *t* test. Expression values are mean ± SEM).

In whole-cell patch-clamp, we measured the ionic currents evoked by NMDA, a specific agonist of NMDARs (Watkins, 1981), in D60 iPSC-derived neurons. To do so, we perfused the cells with a solution containing 100 μM NMDA and 10 μM Glycine, a co-agonist that facilitates the opening of the NMDARs (**Figure 3B**). These conditions aim to saturate the receptors to obtain the highest possible response. The average amplitude of the ionic currents were 342 ± 346 pA and 198 ± 250 pA in cells cultured in BPM and NMM, respectively (**Figure 3C**). The ionic currents also exhibited a similar time to decay to approximately 36.8% of their maximum amplitude (tau of desensitization= 1.4 ± 0.61 s and 0.99 ± 0.61 s in cells cultured in BPM and NMM, respectively, ns: *p*= 0.159; **Figure 3D**). Altogether, these electrophysiological findings demonstrate the expression of functional NMDARs in D60 iPSC-derived neurons.

Our finding of increased expression of *GRIN2A* in BPM condition (Figure 2G), motivated us to evaluate the relative contribution of GluN2B and GluN2A to the NMDA-mediated ionic currents in D60 iPSC-derived neurons. To study the GluN2B contribution we used the selective blocker Ro 25-6981 (Fischer et al., 1997). After NMDA + Glycine perfusion for one second, addition of 1nM Ro 25-6981 resulted in a decrease of ∼30 ± 9% and ∼20 ± 8% of the ionic current of cells cultured in BPM and NMM, respectively (*p*= 0.041) (**Figure 3E, F**). The discrete inhibition of the NMDA-mediated ion currents by Ro 25-6981 suggested that GluN2A could be the main subunit that mediated these currents. Accordingly, perfusion of 300 nM NVP-AAM077, a selective GluN2A blocker (Auberson et al., 2002) resulted in higher values of current inhibition (53 ± 21% and 39 ± 28% in BPM and NMM conditions, respectively, **Supplementary Figure 2**). These results indicate that GluN2A is a critical component of NMDAR-mediated currents in iPSC-derived neurons at D60. This is a fact for cells in BPM conditions, but also for cells in NMM, despite their low mRNA GluN2A expression levels. Overall, these data provide evidence of a mature phenotype in our cultured neurons.

### GluN2A drives changes in intracellular calcium signaling in human glutamatergic neurons

NMDAR activation ultimately increases the intracellular Ca^2+^ concentration in neurons, which is key as secondary messenger for different functions. Thus, an increase in intracellular Ca^2+^ is considered a proxy for evaluating an integrated neuronal response (Brini et al., 2014; Grienberger & Konnerth, 2012).

To evaluate NMDAR-mediated changes in the intracellular Ca^2+^ concentration, the Ca^2+^-sensitive probe Fura-2 was used in the D60 iPSC-derived neural cells, which showed similar basal intracellular Ca^2+^ levels (without external input) in both BPM and in NMM conditions (**Figure 4A**). Perfusion of 100 μM NMDA and 100 μM Glycine induced a change in the Fura-2 signal (ΔF340/380) in 67% of the neurons cultured in BPM, and only in 12% of the cells cultured in NMM conditions (**Figure 4B**). Additionally, the Fura-2 signal induced by NMDA was significantly higher in BPM conditions (0.32 ± 0.18) respect to NMM (0.23 ± 0.10; **Figure 4C**).

**Figure 4.**
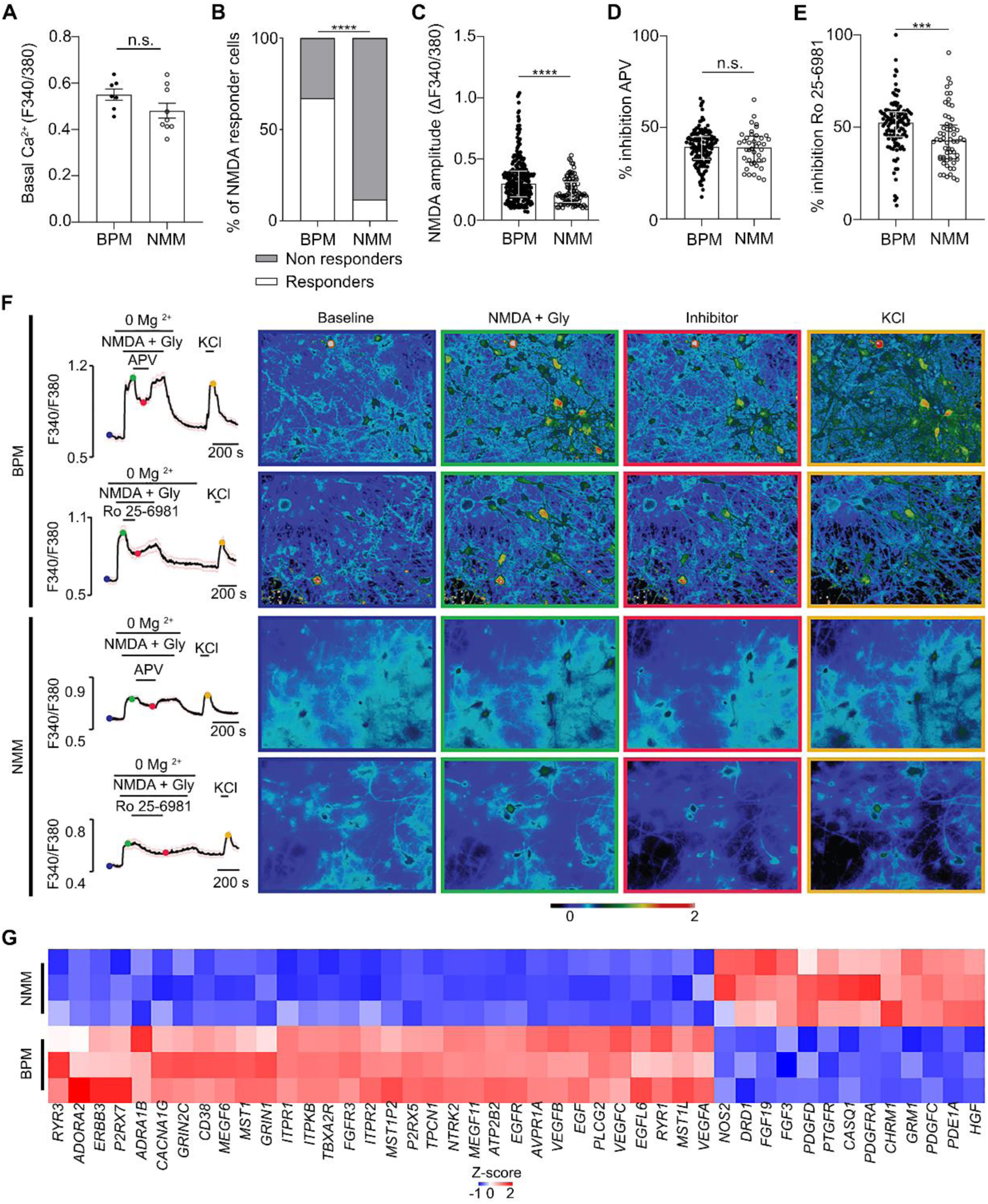
Intracellular calcium changes in iPSC-derived neural cultures in BPM and NMM conditions after 60 days in culture. **A)** Quantification of Fura-2 signal (F340/380) under no stimulation in neural cultures (n= 7 for BPM, n= 9 for NMM). **B)** Percentage of cells that respond to NMDA (n= 399 for BPM, n= 837 for NMM; deviation of > 0.1 in the Fura-2 signal from the baseline, \*\*\*\**p* < 0.0001; Fisher’s exact test). **C)** Amplitude of Fura-2 signal induced by NMDA perfusion (n= 268 for BPM, n= 98 for NMM; \*\*\*\**p* < 0.0001; Mann-Whitney test). **D)** Percentage of Fura-2 signal inhibition by APV respect to NMDA Fura-2 signal (n= 152 for BPM, n= 41 for NMM). **E)** Percentage of Fura-2 signal inhibition induced by Ro-6981 respect to NMDA Fura-2 signal (n= 117 for BPM, n= 58 for NMM; \*\*\**p* < 0.0002; Mann-Whitney test). **F)** Representative average traces of the Fura-2 signal in neural cells cultured in BPM and NMM at basal conditions, after perfusion of the agonist NMDA (without Mg^2+^), the addition of either the NMDAR antagonist APV (n= 52 for BPM, n= 11 for NMM) or the GluN2B inhibitor Ro 25-6981 (n= 33 for BPM and n= 11 for NMM), and finally perfusion of KCl. The pictures at the right side illustrate the Ca^2+^ signal by Fura-2 at different time points indicated by coloured dots on the trace. **G)** Heatmap extracted from the RNA-seq showing the 45 up (red)- and down (blue)-regulated genes in the GO cluster “Calcium signaling”. Expression values are mean ± interquartile range.

We employed different NMDAR inhibitors to ensure that the changes in the intracellular Ca^+2^ levels were NMDAR-activation dependent. Perfusion of 50 μM 2-amino-5-phosphonovaleric acid (APV), a specific antagonist that blocks all NMDARs (Zhu et al., 2016), decreased Fura-2 signal in a similar manner in both BPM and NMM conditions (38 ± 1% and 40 ± 2% respectively, **Figure 4D, F**). Removal of APV in the continuous presence of the agonist NMDA led to the full recovery of the Fura-2 signal, proving it was specific to NMDAR activation. Perfusion with the GluN2B blocker Ro 25-6981 resulted in a 51± 1% drop of the Fura-2 signal in BPM conditions, which was significantly higher than that for NMM (44 ± 2%, p<0.0002 Mann-Whitney; **Figure 4E, F**).

At the end of all experiments, a potassium chloride (KCl) solution was perfused as a control of neuronal activity. Altogether, these results show that NMDA can evoke an integrated neuronal response in D60 iPSC-derived neurons, and that GluN2A is responsible for approximately 50% of the NMDA-mediated intracellular Ca^+2^ signal.

The observation that cells cultured in BPM exhibit a higher calcium-signal in response to NMDA than those in NMM, led us to explore possible explanations by returning to our RNA-seq data (**Figure 4G**). We looked for genes associated with “Intracellular signaling mediator proteins” and we found 45 differentially expressed when comparing both conditions. Most of the top upregulated genes identified corresponded to BPM conditions. Among them, *CACNA1G*, *RYR3* and *ITPR1* encode for voltage-gated Ca^2+^ channels, subunits of Ryanodine receptors and inositol 1,4,5-trisphosphate receptors, respectively. Other genes included *CD38*, which promotes the release of Ca^2+^ from the endoplasmic reticulum (Lee et al., 2022) and *TPCN1*, which is important for releasing Ca^2+^ from lysosomes (Calcraft et al., 2009). Fewer genes related to intracellular calcium were down-regulated, as *PDE1A* and *GRM1*. Overall, these transcriptomic changes indicate that neural cells cultured in BPM are better equipped to increase intracellular Ca^+2^ levels.

## DISCUSSION

In this work, we have generated mature neurons from human iPSCs that exhibit two distinct characteristics. Firstly, at later stages in culture, there was a significant increase in the number of NMDARs present at the synapse, as revealed through confocal microscopy. Secondly, after 60 days of culture, the subunit GluN2A played a significant role in NMDA responses in neurons, consistently mediating ion currents and approximately half of the variations of calcium signaling. Our results not only identified the functional contributions of the subunits GluN2B and GluN2A in mature glutamatergic neurons but also provides insight into how these subunits may be acting during the development of the cerebral cortex.

It is known that the addition of astrocytes to iPSC-derived neurons favours faster and further neuronal maturation (Hedegaard et al., 2020; Klapper et al., 2019) and that BPM results in neural cultures with higher relative number of astrocytes (Bardy et al., 2015; Jackson et al., 2018; Satir et al., 2020). Our neural cells cultured on BPM exhibited a one-to-one neuron-astrocyte ratio like that described in the human brain (von Bartheld et al., 2016). Consequently, the BPM neurons were more mature, as illustrated by RNA-seq analysis showing 12 gene sets with enrichment scores higher than 4 associated to “neuron differentiation”, “neuron spine”, “neuron fate commitment”, “neuron action potential” and “forebrain neuron fate commitment”. The top-upregulated genes (**Supplementary Table 1**) are involved in the differentiation and maturation of neurons (Dong et al., 2012; Dou et al., 2021; Karalay et al., 2011; Lavado et al., 2010; Roussel-Gervais et al., 2023). The upregulation of *BMP4* is a good example, as this gene has been identified as a driver of neuronal differentiation in iPSC-derived neurons (Shi et al., 2007) and the mouse brain (Moon et al., 2009). Using pathway enrichment analysis, we also identified that the GO term “Glutamatergic synapse” was upregulated in cells cultured in BPM.

The incorporation of NMDARs into synapses is a well-established hallmark of neuronal maturation supported by studies in mouse brain slices or primary neuronal cultures (Gladding & Raymond, 2011; Groc et al., 2006; Harris & Pettit, 2007; Ivanov et al., 2006; Rosenmund et al., 1995; Tovar & Westbrook, 1999, 2002). To our knowledge, no previous study has directly assessed the synaptic incorporation of NMDARs during neuronal maturation in human neurons using colocalization studies. In this study, we used confocal immunofluorescence (Escamilla, Sáez-Valero, et al., 2024; Li et al., 2011; Snyder et al., 2005) to demonstrate that there is an increase in the colocalization of the subunit GluN1 and the presynaptic protein Synaptophysin during neurodifferentiation and maturation. This represents the first direct evidence that the differentiation and maturation of human neurons are accompanied by a progressive increase in synaptic NMDARs and support the notion that these dynamic changes can be studied in this cellular model. This is also clinically relevant because the location of NMDARs in the synaptic or extrasynaptic membranes has been associated with several brain disorders, including stroke (Tu et al., 2010; Wu & Tymianski, 2018), Huntington’s disease (Levine et al., 2010; Milnerwood et al., 2010) and Alzheimer’s disease (Escamilla, Badillos, et al., 2024; Parsons & Raymond, 2014; Yu et al., 2023).

NMDA-induced ion currents exhibited similar amplitude and desensitization in both BPM and NMM conditions. Perfusion of Ro 25-6981, a GluN2B inhibitor, reduced a 25% average the NMDA-induced currents also in both conditions, in agreement with transcriptomics results that did not show differences between them. This blockage suggested that the remaining ∼75% current depends on GluN2A again in both conditions. This could be unexpected due to the higher *GRIN2A* expression observed in BPM by RT-qPCR-although not for RNA-seq, that suggests that inhibitors as Ro 25-6981 or NVP should reflect a higher participation of GluN2A in BPM conditions than in NMM. However, different mechanisms may explain the greater participation of GluN2A in NMDA-induced current in NMM conditions, such as epigenetic factors (Corbel et al., 2015; Rodenas-Ruano et al., 2012), interactions with co-agonists (Ferreira et al., 2017), or other signaling pathways (Groc et al., 2007; Matta et al., 2011; Nolt et al., 2011). Nevertheless, our finding suggests a significantly greater dependence on GluN2A than has been reported. Some studies described that only 10% of NMDA-induced ion currents are GluN2A dependent (Neagoe et al., 2018; Rzechorzek et al., 2016; Zhang et al., 2022; Zhang et al., 2016) and others that glutamatergic currents are conducted only by AMPA receptors (Lam et al., 2017; Zhang et al., 2013).

Activation of glutamate receptors increases intracellular Ca^2+^ concentration. In particular, ionotropic glutamate receptors directly mediate Ca^2+^ entry or indirectly activate voltage-gated Ca^2+^ channels; in addition, activation of phospholipase C-coupled metabotropic glutamate receptors mobilizes Ca^2+^ from inositol 1,4,5-trisphosphatesensitive stores (IP3, Nakamura et al., 2000). NMDA receptors mediate substantial Ca^2+^ influx; recording NMDA-evoked increases in Ca^2+^ concentration provides a method to assess NMDAR function. Furthermore, because action potentials induce Ca^2+^ influx via voltage-gated Ca^2+^ channels, the intracellular Ca^2+^ concentration also serves as an index of the neurons-integrated response to activation of excitatory networks (McLeod et al., 1998).

The intracellular Ca^+2^ response to NMDA was higher in BPM conditions (∼70% of cells, ∼10% in NMM), and remarkably, with a higher Ca^+2^ signal magnitude. Data from our RNA-seq analysis indicate that cells cultured in BPM show an increased expression of genes encoding voltage-gated calcium channels, Ryanodine receptors and IP3 receptors. This upregulation could contribute to the increase in intracellular Ca^2+^ concentration upon NMDAR activation, either through Ca^+2^ influx from the extracellular space or the release from intracellular deposits, ultimately resulting in a higher NMDA-induced calcium signal.

Previous studies on iPSC-derived neurons did not report information on the contribution of NMDAR subunits to the NMDA-induced calcium signal (D’Andrea et al., 2024; Galiakberova et al., 2022; Pruunsild et al., 2017; Ruden et al., 2021). Here, Ro 25-6981 assays suggest that GluN2A and GluN2B contribution to the NMDA-induced calcium signal is quite similar, and furthermore, the subunit that predominates differs between conditions. This demonstrates that GluN2A subunit mediates not only an important part of the ionic currents induced by the agonist NMDA but also participate in the increase in intracellular Ca^2+^ levels, reflecting an integrated neuronal response.

Neural cells differentiated in BPM showed an interesting finding. Three of the top six upregulated genes in these cells belong to the locus *DLK1*-*DIO3*, which is under the control of imprinting and epigenetic mechanisms (Hernandez & Stohn, 2018). This reveals an epigenetic regulation of developmental programs crucial for the maturation of glutamatergic neurons. In fact, the downregulation of long non-coding RNAs (lncRNAs) originated from the *DLK1-DIO3* region impairs the differentiation of iPSCs into neurons (Mo et al., 2015). The coding gene *DIO3* is highly expressed in neurons (Salas-Lucia et al., 2024; Salas-Lucia et al., 2023) and plays essential roles in brain function (Hernandez & Stohn, 2018; Salas-Lucia, 2024), and the gene *DLK1* has been associated with the regulation of neurogenesis (Ferrón et al., 2011; Montalbán-Loro et al., 2021; Surmacz et al., 2012). This observation suggests that the *DLK1-DIO3* region may play a key role in the differentiation and maturation of iPSC-derived neurons, and it provides proof of concept that the cellular model described in our study is well-suited to investigate the regulatory functions of this locus during neuronal differentiation.

In conclusion, our results provide compelling evidence that human iPSC-derived neurons exhibit prominent GluN2A-dependent ionic currents and calcium signaling. By integrating bulk transcriptomics and confocal studies with electrophysiology and calcium imaging, we consistently demonstrate that mature neurons, such as those cultured in BPM conditions, display a distinct NMDA-related phenotype. These mature neurons show an increased synaptic localization of NMDARs, higher *GRIN2A* expression, strong GluN2A-dependent ionic currents and GluN2A-integrated neuronal response. This study provides the groundwork for future research on NMDAR-related mechanisms in mature neurons, potentially paving the way for studies on neurodevelopmental and neurodegenerative diseases.

## METHODS

### Cell culture

The iPSC line used in this research was the N1-001iC2 hPSCreg name UGOTSAi002-B, provided by Prof. Henrik Zetterberg (Institute of Neuroscience and Physiology, University of Gothenburg, Sweden). The donor was a male aged 75-79 years old at collection with sporadic Alzheimer’s disease. The iPSC line was maintained in Matrigel (Corning #356234)-coated 6-well plates and fed with mTeSR™1 (StemCell Technologies) at 37°C, 5% CO_2_. Passages were done every 4-5 days (80% confluence) using ReLeSR™ solution (StemCell Technologies). Lines were routinely screened for the absence of Mycoplasma.

iPSCs were detached using Gentle Cell Dissociation Reagent (StemCell Technologies #100-0485) at 80% confluence and differentiated into neural precursor cells (NPCs) using the STEMdiff™ SMADi Neural Induction Kit (StemCell Technologies). Then, NPCs were expanded and matured using STEMdiff™ Neural Progenitor Medium (StemCell Technologies). NPCs were passaged using accutase (Millipore-Sigma #A6964). From NPCs, cells were differentiated into neurons using two alternative protocols: “NMM condition”, based in Neural Maintenance Media (NMM) which yields a neuronal culture with a small percentage of astrocytes, and “BPM condition”, based in BrainPhys Media (BPM), which yields a higher percentage of astrocytes (Bardy et al., 2015; Jackson et al., 2018; Satir et al., 2020). The BPM contained BrainPhys Neuronal Medium (StemCell Technologies #05790), NeuroCult SM1 Neuronal Supplement 2% (StemCell Technologies #05711), N2 Supplement-A 1% (STEMCELL Technologies #07152), Dibutyryl cAMP sodium salt 1 mM (Millipore-Sigma D0627), L-Ascorbic acid 200 nM (Millipore-Sigma #A0278), BDNF 20 ng/mL (Peprotech #450-02), GDNF 20 ng/mL (Peprotech #450-10), and Penicillin/Streptomicin 0.25x (Gibco #15140122). The NMM contained a 1:1 dilution of DMEM/F12+Glutamax (Gibco #31331028) and Neurobasal medium (Gibco #21103049), supplemented with insulin (Gibco #91077C), β-mercaptoethanol 55 μM (Gibco #21985023), MEM Non-essential Amino Acids 0.5x (Gibco #11140050), Sodium Pyruvate 0.5 mM (Gibco #S8636), N2 Supplement 0.5x (Gibco #17502048), B27 Supplement 0.5x (Gibco #17504001), and Penicillin/Streptomycin 0.25x (Gibco #15140122). The coating where NPCs were plated for differentiation consisted of Poly-L-ornithine solution diluted 1:1 in PBS overnight at 4°C (Millipore-Sigma #P4957), followed by 4h at 37°C of Laminin 2020 (Millipore-Sigma #L2020). During the neural induction period, the expression of Nestin and PAX6 was checked as makers of NPCs, and the expression of MAP2 as a marker of mature neurons.

### Immunocytofluorescence

Cells were fixed in 4% paraformaldehyde for 15 min at room temperature (RT) and stored in phosphate buffered saline (PBS) at 4°C until immunostaining was performed. Cells were washed in PBS once and permeabilized using 0.1% Triton X-100 in PBS for 10 min and blocked in 1% bovine serum albumin in PBS for 1 h at RT. Then, cells were incubated with primary antibodies in blocking solution overnight at 4°C. Primary antibodies used were S100β (rabbit, 1:500, Proteintech 15146-1-AP), GFAP (mouse, 1:500, Thermo Fischer MA5-12023), TUJ1 (beta-3-tubulin)(mouse, 1:500, abcam ab14545), MAP2 (chicken, 1:500, Thermo Fischer PA1-10005), Synaptophysin (rabbit, 1:250, Thermo Fischer MA5-14532), PAX6 (rabbit, 1:500, Thermo Fischer 42-6600), Nanog (mouse, 1:500, BioLegend 674002), OCT4 (rabbit, 1:500, Thermo Fischer MA5-14845), GluN1 N-terminal (rabbit, 1:500, Alomone AGP-046).

After three washes with PBS, secondary antibodies (Alexa Fluor^TM^ 647 goat anti-rabbit IgG (H+L), Cy3^TM^ goat anti-mouse IgG (H+L), Alexa Fluor^TM^ 488 goat anti-guinea pig IgG (H+L), Alexa Fluor^TM^ 488 goat anti-chicken IgY (H+L), Thermo Fischer) were added in blocking solution for 1 h RT in the dark. 4’,6- diamidino-2-phenylindole (DAPI, 1 μM) was added as a nuclear counterstain before the last wash with PBS. Cells were mounted on slides using ProLong^TM^ Diamond Antifade Mountant (Invitrogen, P36961). Images were taken by a Leica SPEII confocal microscope and visualized using Imaris.

To assess the synaptic localization of NMDA receptors during neuronal maturation, we performed co-localization analyses between GluN1, the obligatory subunit of NMDARs, and Synaptophysin, a presynaptic vesicle protein. Immunostained iPSC-derived neurons at differentiation days 30 and 60 were imaged using a Leica SPEII confocal microscope under identical acquisition settings to allow for quantitative comparison. Z-stack images were acquired with a step size optimized to meet Nyquist sampling criteria.

For co-localization quantification, raw images were deconvolved using Huygens Professional software (Scientific Volume Imaging, The Netherlands) with a classic maximum likelihood estimation (CMLE) algorithm. Point Spread Functions (PSFs) were automatically generated based on imaging parameters, and signal-to-noise ratios were set to ensure optimal restoration quality. Quantification of co-localization was performed using Mander’s coefficient, which calculates the proportion of one fluorophore’s signal that overlaps with the other within the region of interest. Specifically, Mander’s M1 coefficient was used to determine the fraction of GluN1 signal overlapping with Synaptophysin, thereby estimating the proportion of NMDARs localized at synaptic sites. The regions of interest were defined on TUJ1-positive neurons to restrict the analysis to neurons.

### Fluorescence-activated Cell Sorting (FACS)

Cells were collected using Gentle Cell Dissociation Reagent (StemCell Technologies) for iPSCs and accutase for the neural culture at day 60 to dissociate them into single cells. Cells were counted and collected in 15 mL Falcon tubes, centrifuged at 1000 rpm for 3 min and fixed in PBS 4% for 10 min at RT. Afterwards, cells were centrifuged at 1000 rpm for 3 min and resuspended in PBS for their distribution in Eppendorf tubes (200.000 cells/tube), and centrifuged at the same speed and time before incubation with conjugated antibodies for 30 min at RT in the dark (5 µL conjugated antibodies + 100 µL PBS with Triton X-100 0.01% (w/v) for 4 tubes). Finally, cells were centrifuged and resuspended in PBS until analysis. A negative control was always included, as well as single conjugated antibodies when the assay included more than one simultaneous marker. FACS was performed with BD FACS Aria™ III for iPSCs and BD FACSDiscover™ S8 Cell Sorter for iPSC-derived neural cells at day 60.

### RT-qPCR

RNA was extracted from NPCs, NMM or BPM iPSC-derived neural cells using the PureLink™ Micro Total RNA Purification System (Life Technologies, Carlsbad, CA, USA) following the manufacturer’s instructions. SuperScript™ III Reverse Transcriptase (Life Technologies, Carlsbad, CA, USA) was used to synthesize cDNAs from this total RNA (2 μg) using random primers according to the manufacturer’s instructions. Real time Quantitative PCR (RT-qPCR) amplification was performed on a QuantStudio3 (Applied Biosystems, Thermo Fisher Scientific, Rockford, USA) with Thermo Fisher Scientific TaqMan probes specific for human *GRIN2B* (assay ID: Hs01002012_m1)*, GRIN2A* (assay ID: Hs00168219_m1)*, MAP2* (assay ID: Hs00258900_m1)*, GFAP* (assay ID: Hs00909233_m1), and human *GAPDH* as a housekeeping gene (assay ID: Hs02786624_g1) to normalize the expression levels of the target gene by comparative 2^−ΔCt^ method.

### RNA-seq

RNA was extracted from NMM and BPM iPSC-derived neural cells using the Arcturus^®^ PicoPure^®^ Frozen RNA Isolation Kit (Thermo Fisher) following the manufacturer’s instructions. Samples of total RNA were sent to the genomic facility of the Centre for Genomic Regulation, Barcelona, Spain, for library preparation and sequencing. Only samples of RNA with RNA integrity number (RIN) above 9.5 were analyzed in the RNA-seq. Libraries were paired-end sequenced with a read length of 1x50bp, 30 million reads per sample, and using NextSeq 2000. The FASTQ files were aligned to the Gencode hg38 transcriptome with STAR (v.2.7.8a) using the Partek Flow platform. Aligned reads were quantified to the annotation model (Partek E/M) and normalized to counts per million. A fold change of ≤ 2 or ≥ 2 was applied to filter the data. Only changes with a *p*-value below 0.05 were considered significant.

### Electrophysiological recordings and perfusion procedures

We performed electrophysiology experiments in iPSC-derived neurons cultured in BPM and NMM at D60 of differentiation. Membrane currents elicited by NMDA (TOCRIS, HB0454) and Glycine (Millipore Sigma, 106-57-0) were recorded at a membrane potential of -60 mV using the whole-cell configuration of the patch-clamp technique with a List Electronics EPC-7 amplifier. Patch electrodes were fabricated from borosilicate glass with a resistance of 3-5 MΩ. Currents were filtered at 2 kHz and transferred at a sampled rate of 1-5 kHz into a personal computer for analysis using pCLAMP Acquisition and Analysis (Molecular Device). Experiments were performed at room temperature. iPSC-derived neurons were rapidly perfused using a linear array of glass tubes placed 200-300 mm from the cell body and operated by a motorized device under the control of the computer. Ringer solution containing the different agonists and antagonists flow through adjacent tubes, and solution exchange was achieved by lateral displacement of the whole prefusion array.

Experimental solutions. The normal external solution for iPSC-derived neural cells was (in mM): 140 NaCl, 2.5 KCl, 1.8 CaCl_2_, 15 mM glucose, and 10 mM HEPES at pH 7.4. MgCl_2_ was not included to avoid the voltage-dependent block of NMDARs by Mg ions. Pipettes were filled with this internal solution (in mM): 130 cesium methanesulfonate, 4 NaCl, 0.2 EGTA, 10 HEPES, 10 tetraethylammonium, 2 Na_2_ATP, 0.5 Na_3_GTP, 5 QX314, pH 7.3. Osmolarity was adjusted to 300 and 290 mOsm for external and internal solutions, respectively. Agonists and antagonists were diluted from stock solutions to the desired concentration in external solution: 100 μM NMDA, 10 μM Glycine, 1 μM Ro 25-6981 (TOCRIS, 1594), and 300 nM NVP-AAM077 (MedChemExpress, HY-12294A). To record excitatory postsynaptic current (EPSCs), the external solution was replaced with the following one (in mM): 124 NaCl, 2.5 KCl, 1.25 KH_2_PO_4_, 2 CaCl_2_, 26 NaHCO_3_, and 15 glucose, equilibrated with 95% O_2_/5% CO_2_, pH 7.3 (300 mOsm), supplemented with picrotoxin 100 μM to inhibit IPSC (inhibitory postsynaptic currents).

### Ca^2+^ imaging

Cells were incubated with 5 μM Fura2-AM, a fluorescent probe that changes its wavelength emission from 340 to 380 nm when binds to Ca^2+^, and 0.2% Pluronic F-127 (Thermo Fisher Scientific) in the control external solution for 45 min at 37°C in a 5% CO_2_ incubator. Coverslips with cultured cells were placed in a low-volume chamber continuously perfused at ∼1 ml/min on an inverted microscope (model DMI 3000B, Leica). Fluorescence signals were obtained by exciting Fura2 at 340 and 385 nm with a LED-based system (Lambda 721, Sutter Instruments) and filtering the emitted fluorescence with a 510 nm long-pass filter. Images were acquired using an Orca ER CCD camera (Hamamatsu Photonics) and analyzed with Metafluor software (Molecular Devices). Increases in cytosolic Ca^2+^ are presented as the ratio of the emission intensities at 340 and 380 nm (ΔF340/F380; fluorescence arbitrary units). A positive calcium response was considered when the fluorescence signal (ΔF340/F385) deviated by at least 0.1 of the baselines. Experiments were performed at a temperature of 33–34°C, maintained by a homemade water-cooled Peltier system controlled by a feedback device. NMDA 100 µM + Glycine 100 µM (Millipore Sigma, 106-57-0) was applied extracellularly.

The control external solution contained the following (in mM): 140 NaCl, 2.5 KCl, 1.8 CaCl_2_, 1 MgCl_2_, 15 glucose, and 10 HEPES (300 mOsm/kg adjusted to pH 7.3 with NaOH). Free-Mg^2+^ solution contained the following (in mM): 140 NaCl, 2.5 KCl, 1.8 CaCl_2_, 15 glucose, and 10 HEPES (298 mOsm/kg adjusted to pH 7.3 with NaOH). KCl (50mM) solution contained the following (in mM): 9.2 NaCl, 50 KCl, 1.8 CaCl_2_, 1 MgCl_2_, 15 glucose, and 10 HEPES (298 mOsm/kg adjusted to pH 7.3 with NaOH). NMDA 100 μM + Glycine 100 μM was applied extracellularly in free Mg^2+^ solution. NMDA receptor blockers: APV (TOCRIS, 0106) 50 μM, and Ro 25-6981 1 μM, were added to NMDA 100 μM + Glycine 100 μM in free MgCl_2_ solution.

The OriginPro 2020 software (OriginLab) was used for graphing. 3-5 experiments were performed in each condition from cultured cells of 2-3 different days.

### Statistical analysis

The distribution of data has been tested for normality using a D’Agostino-Pearson test. ANOVA has been used for parametric variables, and the Kruskal-Wallis test for non-parametric variables for comparison between groups. A Student’s t-test for parametric variables and a Mann-Whitney U test for non-parametric variables were employed to compare the two groups and determine *p*-values. A Z-test was used for analyzing calcium imaging data. When considered, the results are presented as the means ± SD or means ± SEM (see figure legends), and all the analyses were performed using GraphPad Prism (version 7; GraphPad Software, Inc).

## Acknowledgements

We thank Prof. Juan Lerma for generously providing access to his laboratory facilities for the patch clamp experiments, for his assistance with data analysis and for his insightful contributions that greatly enriched this work.

---

**Supplementary Figure 1.**
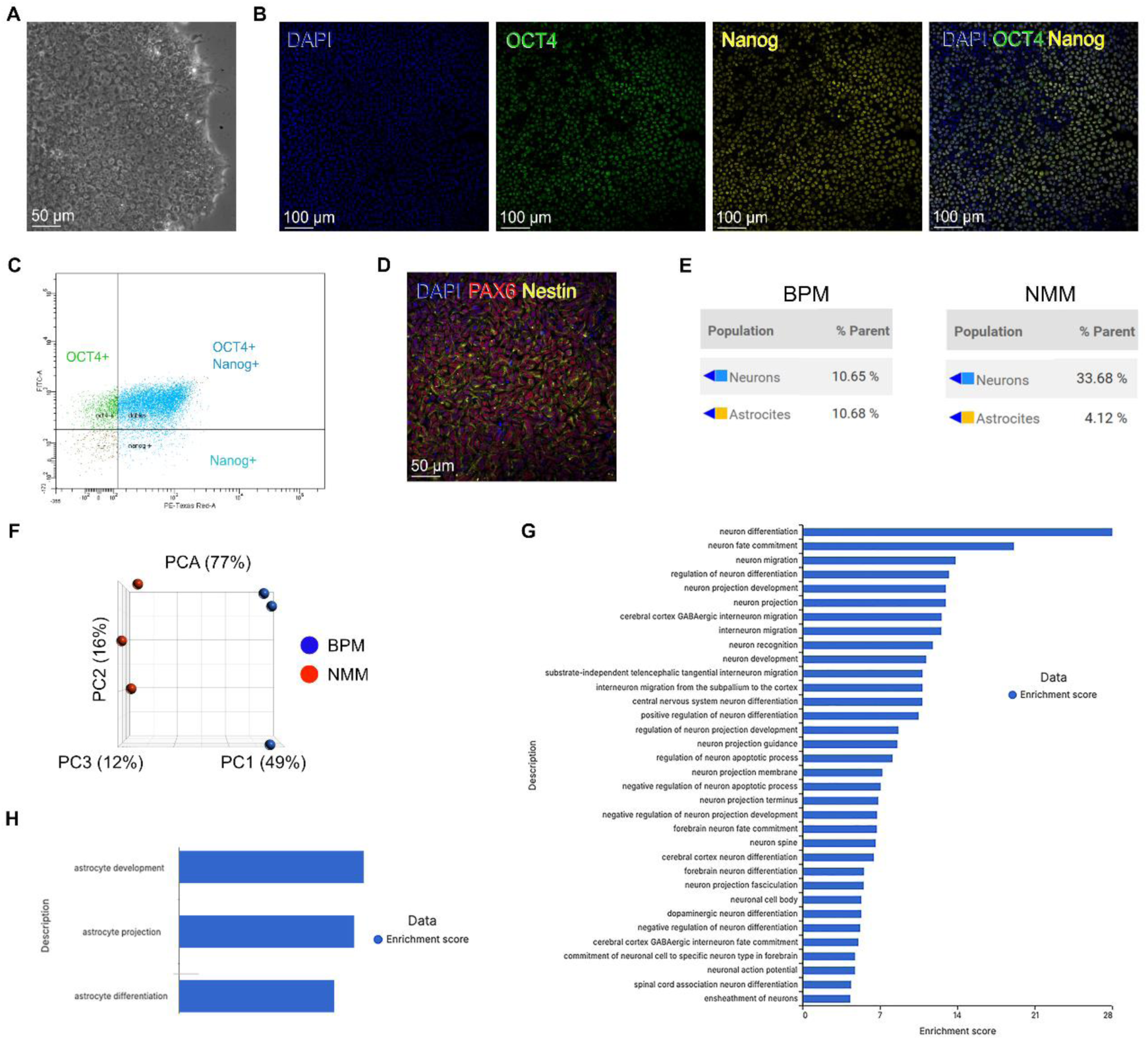
**A)** Brightfield image showing a compact iPSCs colony. **B)** Immunofluorescence images of iPSCs showing staining against the pluripotency markers OCT4, Nanog and DAPI. **C)** Dispersion plots from a fluorescence-activated cell sorting (FACS) experiment showing cell singlets. The OCT4 antibody was conjugated to an AlexaFluor 488 fluorophore, while the Nanog antibody was linked to an AlexaFluor 594 fluorophore. Doublets corresponding to cells expressing both markers were > 80%. **D)** Immunofluorescence image of neural precursor cells (NPCs) showing staining against PAX6, Nestin and DAPI. **E)** Quantification of singlets within the black boxes of Figure 1D. **F)** Principal component plot extracted from RNA-seq data illustrating differences between iPSC-derived neurons cultured in BPM and in NMM at day 60 of differentiation. **G)** List of Gene Ontology (GO) terms enriched in cells cultured in BPM with respect to NMM containing the keyword “neuron”. **H)** List of GO terms enriched in cells cultured in BPM with respect to NMM containing the keyword “astrocyte”.

**Supplementary Figure 2.**
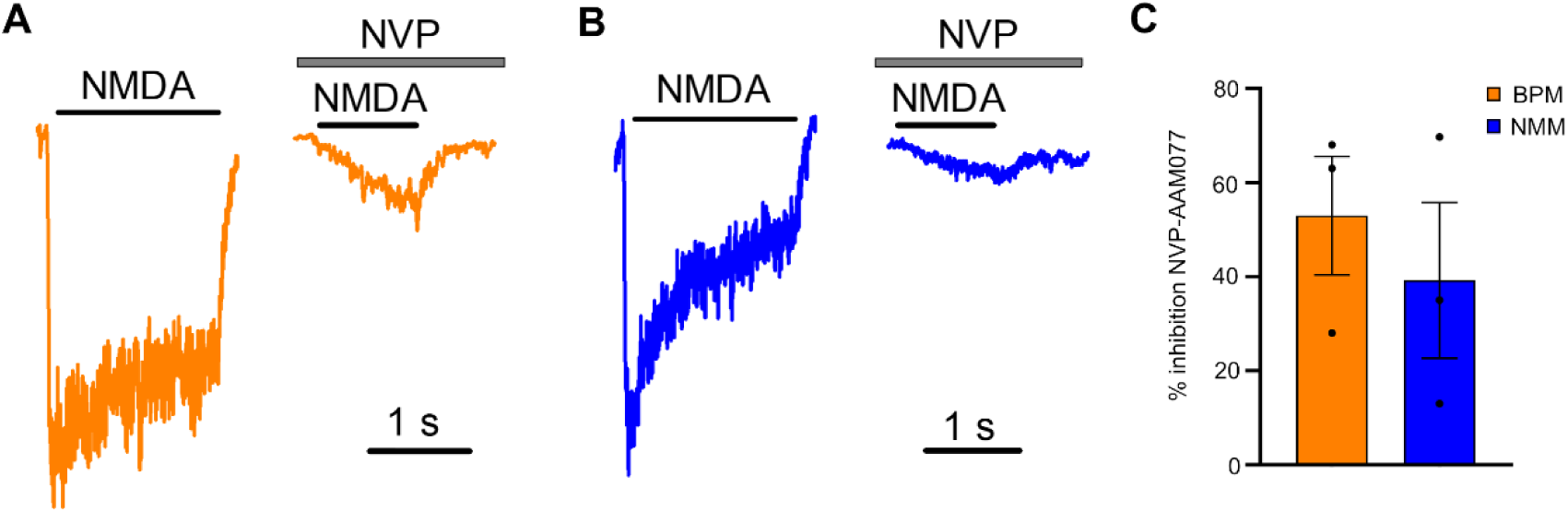
Representative whole-cell recording of membrane currents upon NMDA perfusion (left) and after addition of the GluN2A antagonist NVP-AAM077 (right) on the same cell, in iPSC-derived neurons cultured in BPM (**A**) or in NMM (**B**) conditions. **C)** Quantification of GluN2A-current inhibition by NVP-AAM077 (n= 3 for BPM, n= 3 for NMM) Expression values are mean ± SEM.

